## Supplementary Information for "A *MSTN*^Del273C^ mutation with *FGF5* knockout sheep by CRISPR/Cas9 promotes skeletal muscle myofiber hyperplasia"

### Table S1 Primers sequences of gene cloning

| Genes | Primer Name | Primer Sequences（5’-3’） |
| --- | --- | --- |
| FOSL1 | pFOSL1-F | tttgccgccagaacacaggaccggttctagaGCCACC**atgttccgagactacggggaac** |
|  | pFOSL1-R | agagagaagtttgttgcgccggatcccttatcgtcatcgtctttgtaatc**taaggccaagagggttggggag** |

Note: The bold is the FOSL1 gene sequences, the Kozak sequence in the box, the underline position is the flag tag sequence, and the lowercase letters are the homologous arm sequence of the vector.

### Table S2 The sequences of siRNA

| Sequences Name | Sequences (5’-3’) |
| --- | --- |
| si-oar-FOSL1_001 | GGAAAGAACTGACCGACTT |
| si-oar-FOSL1_002 | AACCCTCTTGGCCTTATGA |
| si-oar-FOSL1_003 | CCAAGCATCAACACTGTGA |

### Table S3 All primers of PCR and RT-qPCR

| Genes |  | Primer sequences（5’-3’） | Product length (bp) |
| --- | --- | --- | --- |
| MSTN | F  R | AAGTCAAGGTAACAGACACACC  CAATACTACATATAGATTTTTC | 756 |
| FGF5 | F  R | ATTTCAAGAAAACAGCTATAAT  AAAAGTTGCCTTCAGAGCACT | 396 |
| GAPDH | F | GTCGGAGTGAACGGATTTGG | 97 |
|  | R | TGAAGGGGTCATTGATGGCA |  |
| MSTNdel | F | CAATTACTGCTCTGGAGAATGTG | 121 |
|  | R | AGACATCTTTGTAGGAGTACAGC |  |
| Cyclin A1 | F | CCAAGGCACACTACATGAGGAA | 133 |
|  | R | AGGAAGTTGACAGCCAGGTAGA |  |
| Cyclin B1 | F | TGACTGACAACACCTACACCAA | 142 |
|  | R | AGCTCAACATCAACCTCTCCAA |  |
| Cyclin D1 | F | CATCGAGCACTTCCTCTCCAA | 111 |
|  | R | GAAATGAACTTCACGTCTGTGGC |  |
| Cyclin E1 | F | CAGTGTGGCAGTCAGCCTTG | 123 |
|  | R | CCTCTGAGGCTTGTACGCAG |  |
| CDK1 | F | CATGAGGTAGTGACACTCTGGTA | 127 |
|  | R | ACAGTGGTTTCTTCGTTGCTAA |  |
| CDK2 | F | TACCCCATACCCCGTGACCC | 108 |
|  | R | GGCCAAACCACCTCATCTGG |  |
| CDK4 | F | ACATTCTGGTGACAAGTGGTGG | 85 |
|  | R | GGTGTAAGTGCCATCTGGTAGC |  |
| CDK6 | F | GCCTGGACTTTCTTCATTCTCAC | 144 |
|  | R | CCACTGAGGTAAGAGCCATCTG |  |
| PCNA | F | CAAGTGGCGTGAACCTACAG | 142 |
|  | R | AGTATTTTGGACATGCTGGTGAG |  |
| BCL2 | F | TTGGGAAGTTTTCAGAGCAGC | 140 |
|  | R | CCTCCTCCGTGATGTGGTAT |  |
| MKI67 | F | CTGGTGTCAAGAGTCGGCTAAGA | 141 |
|  | R | GGCAGGACGCTGGAGTGATT |  |
| MyoD1 | F | CCCCAACCCGATTTACCAGG | 119 |
|  | R | TAAGCGCAATCTTTTGGGCG |  |
| MyoG | F | CCAGTGAATGCAGCTCCCATA | 132 |
|  | R | AGGTGAGGGAGTGCAGATTG |  |
| MyHC | F | ATGAGGGGGACACTGGAAGA | 113 |
|  | R | CGGATGAACTTGCCAAAGCG |  |
| FOSL1 | F | GGTGTTTCTGATGCTCGCTG | 113 |
|  | R | GGCTAGAGCTTGATGCGGTT |  |
| c-Fos | F | AAGGAGAATCCGAAGGGAAAGG | 100 |
|  | R | TAGTTGGTCTGTCTCCGCTT |  |
| cMyoD1-1 | F | GACGGACTGCAAGGAGGAAG | 147 |
|  | R | GCTCAGATCGCTCCAATATCCA |  |
| cMyoD1-2 | F | CATCGTCACAGGTGGCATCA | 90 |
|  | R | TCTCCAGCACCCTGACTAAATC |  |
| Rac1 | F | ACCCGCAGACAGATGTATTCT | 139 |
|  | R | TCAAGTTTCGTCCCCACCAG |  |
| RYR1 | F | CCACAACTTTAAGCGCGAGG | 113 |
|  | R | ACTGTACATCTCCCGCCTTG |  |
| RYR3 | F | AAAGTATGGGCCCGAAGTGG | 107 |
|  | R | CCGCCTGTGCTTTCCTTTTC |  |
| MYMK | F | CCAGAAGGCGGTTCCACAT | 146 |
|  | R | CGTCCCGTAGATGCTGAAGTA |  |
| MYMX | F | GCTGTCTGTTGTTCGTCCTCA | 90 |
|  | R | CTAGCCTCTCCCTCCTTTCCA |  |
| DMD | F | TGCACTACTCTCAACAGCGT | 131 |
|  | R | GGCTGAAAGGAGCCATGAGA |  |
| SMN1 | F | GGTGTAGCTCTCTCCCAAGGA | 109 |
|  | R | ATGCAGATGACATGCACCCA |  |
| MTM1 | F | CCAGTGGAGCAGCGTTACAT | 110 |
|  | R | GAGGCCGAGGGATCAGAAAG |  |
| IGF1 | F | TCACATCCTCCTCGCATCTCTT | 130 |
|  | R | CTGTCTCCGCACACGAACTG |  |
| GAA | F | CAGCCGCAGGAACCATACAG | 107 |
|  | R | CCGTGGAACAGCGTGTAGAG |  |
| ACVR1 | F | TGCCTTCTAGCCTGCCTACTG | 117 |
|  | R | GTTGGTGGTGATGAGCCCTTC |  |
| ACVR2A | F | CCAGTAACACCTAAGCCTCCC | 142 |
|  | R | GAGTTGGAACGAGCACAGGA |  |
| ACVR2B | F | CCAGCTCTGCGTGACAATC | 107 |
|  | R | CGTTGACCGACCTCCGAAT |  |

### Table S4 The antibodies information

| Antibody Name | Manufacturer | Catalog Number | Host |
| --- | --- | --- | --- |
| GAPDH | Zhongshan Golden Bridge | TA-08 | Mouse |
| β-Tubulin | Zhongshan Golden Bridge | TA-10 | Mouse |
| MyoD1 | Affinity | AF7733 | Rabbit |
| MyoG | DSHB | F5D | Mouse |
| MyHC | DSHB | MF20 | Mouse |
| FOSL1 | Affinity | AF6921 | Rabbit |
| p-FOSL1 | Affinity | AF7235 | Rabbit |
| MEK1/2 | Affinity | AF6385 | Rabbit |
| p-MEK1 | Beyotime | AF1786 | Rabbit |
| ERK1/2 | Affinity | 4695 | Rabbit |
| p-ERK1/2 | Affinity | 4370 | Rabbit |
| p38 MAPK | Affinity | AF6456 | Rabbit |
| p-p38 MAPK | Affinity | AF4001 | Rabbit |
| Pax7 | DSHB | - | Mouse |
| Myosin VIIa | abcam | Ab3481 | Rabbit |
| MSTN | Affinity | DF13273 | Rabbit |

### Table S5 Summary of generation of sheep carrying biallelic mutations in dual genes via the CRISPR/Cas9 system

| Cas9/sgRNA  (molar ratio) | Embryo injected | Recipients | Pregnancy | Alive lambs | Mutation | | | Biallelic mutation | | |
| --- | --- | --- | --- | --- | --- | --- | --- | --- | --- | --- |
|  |  |  |  |  | MSTN | FGF5 | MSTN+FGF5 | MSTN | FGF5 | MSTN+FGF5 |
| 1:2 | 448 | 93 | 35 | 28 | 1 | 3 | 1 | 0 | 0 | 0 |
| 1:10 | 365 | 84 | 26 | 22 | 2 | 2 | 2 | 2 | 2 | 2 |
| 1:15 | 388 | 59 | 17 | 14 | 1 | 0 | 0 | 1 | 0 | 0 |

### Table S6 Muscle weight of different parts in WT and MF^+/-^ sheep (g)

| **Muscle Classification** | **WT (n=3)** | **MF^+/-^ (n=4)** | ***P*-value** |
| --- | --- | --- | --- |
| Longissimus dorsi | 905.67±37.15 | 760.00±48.68 | 0.077 |
| Biceps brachii | 50.67±2.67 | 48.25±3.2 | 0.606 |
| Triceps brachii | 247.33±18.41 | 265.00±8.01 | 0.374 |
| Gluteus medius | 305.00±25.24 | 339.75±20.1 | 0.324 |
| Semimembranous | 199.33±21.84 | 194.00±18.5 | 0.859 |
| Semitendinosus | 435.33±41.73 | 400.00±19.71 | 0.438 |
| Biceps femoris | 712.00±57.65 | 619.50±78.39 | 0.417 |
| Quadriceps femoris | 604.67±46.23 | 612.50±29.5 | 0.886 |

### Table S7 The slaughter traits of muscles in WT and MF^+/-^ sheep

| **Slaughter Indexes** | **WT (n=3)** | **MF^+/-^ (n=4)** | ***P*-value** |
| --- | --- | --- | --- |
| Live weight (kg) | 56.33±3.088 | 50.15±2.058 | 0.14201 |
| Carcass weight(kg) | 32.23±2.436 | 28.5±1.588 | 0.23588 |
| Slaughter percentage (%) | 57.12±1.237 | 56.75±1.259 | 0.84403 |
| loin muscle area (cm^2^) | 17.17±1.58 | 13.95±1.757 | 0.24795 |
| Meat weight (kg) | 18.79±1.306 | 15.68±0.825 | 0.08707 |
| The proportion of meat in carcass | 0.58±0.005 | 0.55±0.018 | 0.18691 |
| The proportion of brisket and neck meat | 0.14±0.018 | 0.13±0.004 | 0.85322 |
| The proportion of loin meat | 0.11±0.005 | 0.09±0.011 | 0.33339 |
| The proportion of rib meat | 0.22±0.004 | 0.15±0.003 | 0.00003 |
| The proportion of foreleg meat | 0.18±0.009 | 0.21±0.012 | 0.20974 |
| The proportion of hind leg meat | 0.33±0.009 | 0.4±0.016 | 0.0252 |
| Neat percentage (%) | 0.58±0.005 | 0.55±0.018 | 0.18691 |

### Table S8 Meat quality of longissimus dorsi in WT and MF^+/-^ sheep

| **Meat Quality** | **WT (n=3)** | **MF^+/-^ (n=4)** | ***P*-value** |
| --- | --- | --- | --- |
| pH_45min_ | 6.37±0.136 | 6.38±0.043 | 0.95934 |
| pH_24h_ | 5.58±0.038 | 5.49±0.038 | 0.230 |
| L* | 32.14±2.165 | 29.81±2.165 | 0.326 |
| a* | 13.44±0.452 | 15.29±0.452 | 0.084 |
| b* | 7.52±0.990 | 6.22±0.990 | 0.224 |
| Drip loss (%) | 24.84±1.802 | 26.11±1.802 | 0.446 |
| Cooking loss (%) | 30.62±0.864 | 29.15±0.864 | 0.314 |

### Table S9 Shearing force of different parts in WT and MF^+/-^ sheep (N)

| **Muscle Classification** | **WT (n=3)** | **MF^+/-^ (n=4)** | ***P*-value** |
| --- | --- | --- | --- |
| Longissimus dorsi | 42.7±5.497 | 40.5±6.080 | 0.808 |
| Biceps brachii | 37.7±2.125 | 40.2±8.675 | 0.806 |
| Triceps brachii | 49.8±7.174 | 55.5±5.023 | 0.530 |
| Gluteus medius | 53.2±4.946 | 68.0±7.206 | 0.181 |
| Semimembranous | 54.2±3.786 | 64.3±8.376 | 0.379 |
| Semitendinosus | 49.6±4.476 | 45.1±6.445 | 0.616 |
| Biceps femoris | 37.3±6.467 | 54.1±4.721 | 0.084 |
| Quadriceps femoris | 47.2±3139 | 57.0±7.298 | 0.325 |

### Table S10 Amino acid content of longissimus dorsi in WT and MF^+/-^ sheep (%, DM basis)

| **Amino Acid** | **WT (n=3)** | **MF^+/-^ (n=4)** | ***P*-value** |
| --- | --- | --- | --- |
| Aspartic acid (Asp) | 5.62±0.449 | 6.22±0.348 | 0.334 |
| Threonine (Thr) | 2.68±0.211 | 3.00±0.162 | 0.271 |
| Serine (Ser) | 2.11±0.164 | 2.40±0.133 | 0.222 |
| Glutamic acid (Glu) | 10.71±0.848 | 12.18±0.61 | 0.207 |
| Proline (Pro) | 2.04±0.164 | 2.33±0.134 | 0.224 |
| Glycine (Gly) | 2.62±0.203 | 2.92±0.171 | 0.311 |
| Alanine (Ala) | 3.47±0.272 | 3.87±0.210 | 0.285 |
| Cystine (Cys) | 0.54±0.050 | 0.57±0.029 | 0.707 |
| Valine (Val) | 3.20±0.256 | 3.52±0.196 | 0.356 |
| Methionine (Met) | 1.70±0.139 | 1.87±0.101 | 0.353 |
| Isoleucine (Ile) | 3.03±0.246 | 3.38±0.183 | 0.295 |
| Leucine (Leu) | 5.09±0.406 | 5.62±0.318 | 0.346 |
| Tyrosine (Tyr) | 2.11±0.186 | 2.06±0.338 | 0.912 |
| Phenylalanine (Phe) | 3.15±0.279 | 3.48±0.207 | 0.376 |
| Lysine (Lys) | 5.32±0.437 | 5.97±0.358 | 0.297 |
| Histidine (His) | 2.22±0.200 | 2.38±0.179 | 0.577 |
